# The small protein SbtC is a functional component of the CO_2_ concentrating mechanism of cyanobacteria

**DOI:** 10.1101/2025.11.25.690420

**Authors:** Peter Walke, Carolin Poppitz, Niklas Menz, Wolfgang R. Hess, Robert Burnap, Martin Hagemann, Stephan Klähn

## Abstract

Oxygenic phototrophs fix CO_2_ via the enzyme ribulose-1,5-bisphosphate carboxylase/oxygenase (RubisCO) that shows relatively low CO_2_ affinity and specificity. To circumvent low and fluctuating CO_2_ concentrations in aquatic systems, cyanobacteria and algae have evolved sophisticated inorganic carbon (Ci) concentrating mechanisms (CCMs). Bicarbonate transporters such as SbtA play a crucial role in the cyanobacterial CCM and hence are tightly regulated at multiple layers. The control of *sbtA* gene expression and corresponding transporter activity involves the PII-like protein SbtB, whose gene is frequently co-transcribed with *sbtA*. Here we report on the discovery of a so far non-annotated gene in the model *Synechocystis* sp. PCC 6803, which is located upstream of the *sbtAB* operon and encodes the small protein SbtC, composed of 80 amino acids. Presence of SbtC was confirmed by immunoblotting after fusing the *sbtC*-coding sequence to a Flag-tag. Similar to *sbtAB*, transcription of the *sbtC* locus is induced by low CO_2_ availability but controlled independently. Mutation of the *sbtC* locus in a wild-type background showed only a mild phenotype even under low CO_2_, but the diurnal growth was impaired as found before in the mutant Δ*sbtB*. Biochemical analysis provided evidence for a trimeric SbtABC complex in the membrane. Recombinant *Synechocystis* strains harboring only SbtA as single Ci uptake system with either deleted *sbtB* or *sbtC* genes showed that bicarbonate leakage from the cell was strongly elevated in both mutants. Our results provide evidence that SbtC contributes to the formation of the SbtAB complex, thereby regulating bicarbonate exchange at the cytoplasmic membrane. Well-conserved SbtC-like proteins encoded in the neighborhood of *sbtAB* exist in many cyanobacterial genomes pointing towards an important role in the cyanobacterial CCM.

## Introduction

Carbon dioxide (CO_2_) is the primary inorganic carbon (Ci) source for autotrophic organisms. These organisms are at the basis of food chains, because primary production via biological CO_2_ fixation is the foundation for the formation of organic matter and biomass at the global scale. In oxygenic photoautotrophs such as cyanobacteria, algae and plants, CO_2_ is fixed by the enzyme ribulose-1,5-bisphosphate carboxylase/oxygenase (RubisCO), which catalyzes the carboxylation of ribulose-1,5-bisphosphate, generating two molecules of 3-phosphoglycerate (3PGA). Although this reaction is crucial for the global carbon cycle, RubisCO is one of the slowest enzymes, catalyzing the fixation of only three to ten molecules CO_2_ per second per molecule of protein (Ellis 2010). Cyanobacterial RubisCO, in particular, has remained adapted to the ancient atmosphere in which CO_2_ concentrations far exceeded those of oxygen (O_2_), resulting in relatively low substrate specificity for CO_2_. Thus, under modern conditions RubisCO frequently reacts with O_2_ instead of CO_2_. This oxygenation reaction leads to the collateral production of 2-phosphoglycolate (2PG), which is toxic to the cell as it inhibits several enzymes of the Calvin-Benson-Bassam (CBB) cycle, including triose-phosphate isomerase and sedoheptulose-1,7-bisphosphatase (Flügel *et al*. 2017). Hence, 2PG must be salvaged by conversion into 3PGA during photorespiration, a multistep process that is energy-consuming and releases CO_2_ (Bowes, Ogren and Hageman 1971; Hagemann *et al*. 2013). Consistently, the carboxylation reaction is regarded as the rate-limiting step of photosynthesis (Hudson *et al*. 1992; Von Caemmerer *et al*. 1997; Spreitzer and Salvucci 2002).

RubisCO’s limited catalytic efficiency under low atmospheric CO_2_ concentrations, combined with constant fluctuations in Ci availability caused by temperature and pH changes in aquatic environments, created evolutionary pressure to prevent oxygenation activity and increase catalytic efficiency for carboxylation. As a result, several phylogenetically independent Ci-concentrating mechanisms (CCMs) have evolved to enrich CO_2_ in the immediate environment of RubisCO (Raven, Cockell and De La Rocha 2008). An efficient CCM also evolved among cyanobacteria, the only prokaryotes that perform oxygenic photosynthesis. Cyanobacteria contribute significantly to the annual CO_2_ fixation and hence, play major roles in global biogeochemical cycles (Rousseaux and Gregg 2014). The cyanobacterial CCM comprises several Ci uptake systems that enable the accumulation of high intracellular concentrations of bicarbonate (HCO_3_^-^) (Price *et al*. 2008; Hagemann, Song and Brouwer 2021). HCO_3_^-^ subsequently diffuses into carboxysomes, prokaryotic microcompartments, where RubisCO is co-localized with carbonic anhydrase, which catalyzes the conversion of HCO_3_^-^ into saturating amounts of CO_2_ (Yeates *et al*. 2008). In cyanobacteria, five distinct Ci transport systems have been described. SbtA and BicA are single-component Na^+^-dependent symporters for HCO_3_ ^-^ (Shibata *et al*. 2002; Price *et al*. 2004, 2008) while the other three are multicomponent complexes. From these, BCT1 belongs to the ABC transporter family with high affinity to HCO_3_ ^-^ (Omata *et al*. 1999), while the two CO_2_ hydrating systems NDH-1_3_ and NDH-1_4_ are specialized forms of the NADPH dehydrogenase complex NDH-1 (Ogawa 1991; Battchikova, Eisenhut and Aro 2011; Schuller *et al*. 2020).

In past years, the **<u>s</u>**odium-dependent **<u>b</u>**icarbonate **t**ransporter **A** (SbtA) received special attention. SbtA was found to form a homotrimer, with each subunit consisting of ten highly conserved transmembrane helices (Fang *et al*. 2021; Liu *et al*. 2021). SbtA shows remarkably high bicarbonate affinity, which makes it the prime candidate for expression in transgenic plants with the aim of supporting crop plant CO_2_ fixation via the cyanobacterial CCM (Du *et al*. 2014; Long *et al*. 2016). In model cyanobacteria such as *Synechocystis* sp. PCC 6803 (hereafter *Synechocystis*), the corresponding *sbtA* gene is clearly responding to different Ci availability (Wang, Postier and Burnap 2004). In particular, *sbtA* and several other genes were found maximally expressed under Ci-limiting conditions (e.g. ambient air), but only show basal expression at elevated CO_2_ levels (>1 % [v/v] CO_2_). This is mainly controlled by the global LysR-type transcriptional repressor NdhR (Figge *et al*. 2001; Omata *et al*. 2001). DNA-binding of NdhR upstream of *sbtA* is determined by the effector molecules 2-oxoglutarate and 2PG, whose intracellular abundances fluctuate in response to Ci status (Daley *et al*. 2012; Jiang *et al*. 2018). NdhR plays a crucial role not only in controlling *sbtA* expression, but also in regulating the transcription of other Ci transporter genes, such as *bicA* and those encoding the NDH-1_3_ complex (Wang, Postier and Burnap 2004; Klähn *et al*. 2015). Furthermore, two other LysR-type transcriptional regulators, CmpR and RbcR, contribute to the activation of genes encoding either components of the CCM or RubisCO itself (Omata *et al*. 2001; Bolay *et al*. 2022). In addition, we showed that cyAbrB2 is involved in Ci-mediated gene expression control as a supplementary regulator of NdhR and CmpR (Orf *et al*. 2016).

The *sbtA* gene is widespread among cyanobacteria and has also been found in several other bacteria (Shibata *et al*. 2002). In many cyanobacteria, including *Synechocystis,* it is located adjacent to another gene, typically co-transcribed with *sbtA* and was therefore annotated as *sbtB*. The function of SbtB remained enigmatic for a long time, but it has since been identified as an important regulator of cyanobacterial Ci acclimation (Selim *et al*. 2018). Based on structural features, SbtB is now considered as a member of the PII superfamily. PII signal transduction proteins are known to interact with various transporters, thereby fine-tuning their activity and balancing carbon and nitrogen metabolism in response to environmental changes (Forchhammer 2004). A similar function is suspected for the SbtB-SbtA pair. The heterologous co-expression of *sbtB* and *sbtA* in *E. coli* suggested a direct inhibitory effect of SbtB on SbtA-mediated HCO_3_ ^-^ transport activity (Du *et al*. 2014), a finding that has recently been supported by the identification of structural interactions (Fang *et al*. 2021). Moreover, *Synechocystis* cells harboring *sbtB* mutations were found to exhibit impaired Ci acclimation, pointing to a broader regulatory function of this PII-like protein (Selim *et al*. 2018). This finding has gained further support from the results of transcriptomic and metabolomic studies (Mantovani *et al*. 2022). SbtB features a characteristic homotrimeric, ferredoxin-like fold and the distinctive T-loop structure, which is required for interacting with its target protein. Like other PII proteins, SbtB can bind various effector molecules, primarily adenyl nucleotides such ATP, ADP, or AMP, but also the second messengers cAMP and c-di-AMP (Selim *et al*. 2018, 2021). Effector binding induces conformational changes in the T-loop structure, thereby modulating the accessibility of target binding sites. The second messenger cAMP has previously been proposed to act as a Ci-sensing signal, since the activity of the cAMP-producing soluble adenylate cyclase is modulated by different Ci concentrations (Hammer, Hodgson and Cann 2006). The binding of the second messenger c-di-AMP has also been linked to glycogen metabolism during diurnal cycles (Selim *et al*. 2021).

In the present study, we report on the discovery of a new player in the CCM-related SbtA/B network. Directly upstream of their genes, we discovered a previously non-annotated gene encoding a well-conserved small protein called SbtC, which is highly Ci responsive mirroring *sbtAB* regulation. Mutations in the *sbtC* gene in wild-type or mutant backgrounds revealed that SbtC contributes to the regulation of SbtA-mediated responses in the Ci acclimation via interactions with SbtB.

## Results

### A non-annotated open reading frame is present upstream of sbtAB in Synechocystis and many other cyanobacteria

With 1120 bp, the intergenic region between *sbtAB* (*slr1512-13*) and the upstream gene *psbP2* (*sll1418*) is unusually long. Moreover, bacterial genomes are typically compact and hence, we assumed that this region potentially harbors genetic information that has not been captured yet. First hints that it is indeed transcribed were obtained through RNA-seq-type transcriptomic analyses of *Synechocystis* that also revealed a high number of previously unknown protein-coding genes with intriguing expression patterns (Mitschke *et al*. 2011; Kopf *et al*. 2014; Spät *et al*. 2023). Accordingly, a transcriptional unit (TU) originating from a specific transcriptional start site and partially covering that region was mapped (Kopf *et al*., 2014, given as TU1641 there). The corresponding TU1641 includes an open reading frame (ORF) that could encode an 80 amino acid protein (**Fig. 1A**). By using the TBLASTN algorithm (Gertz *et al*. 2006), sequences similar to *sbtC* were also identified in other cyanobacteria. In many cases, a similar genetic organization was found (**Fig. 1B**), indicating that this ORF might be associated with the *sbtAB* genes. Given its conserved neighborhood with the *sbtAB* operon, known to be involved in bicarbonate uptake and regulation, we named the new gene *sbtC*. Alignment of the sequences revealed that the central region of the SbtC protein is highly conserved across different cyanobacterial species (**Fig. 1C**). An AlphaFold structure prediction of the protein suggested that these highly conserved regions form two β-sheets and a transmembrane helix, indicating a potential functional relevance.

**Figure 1:**
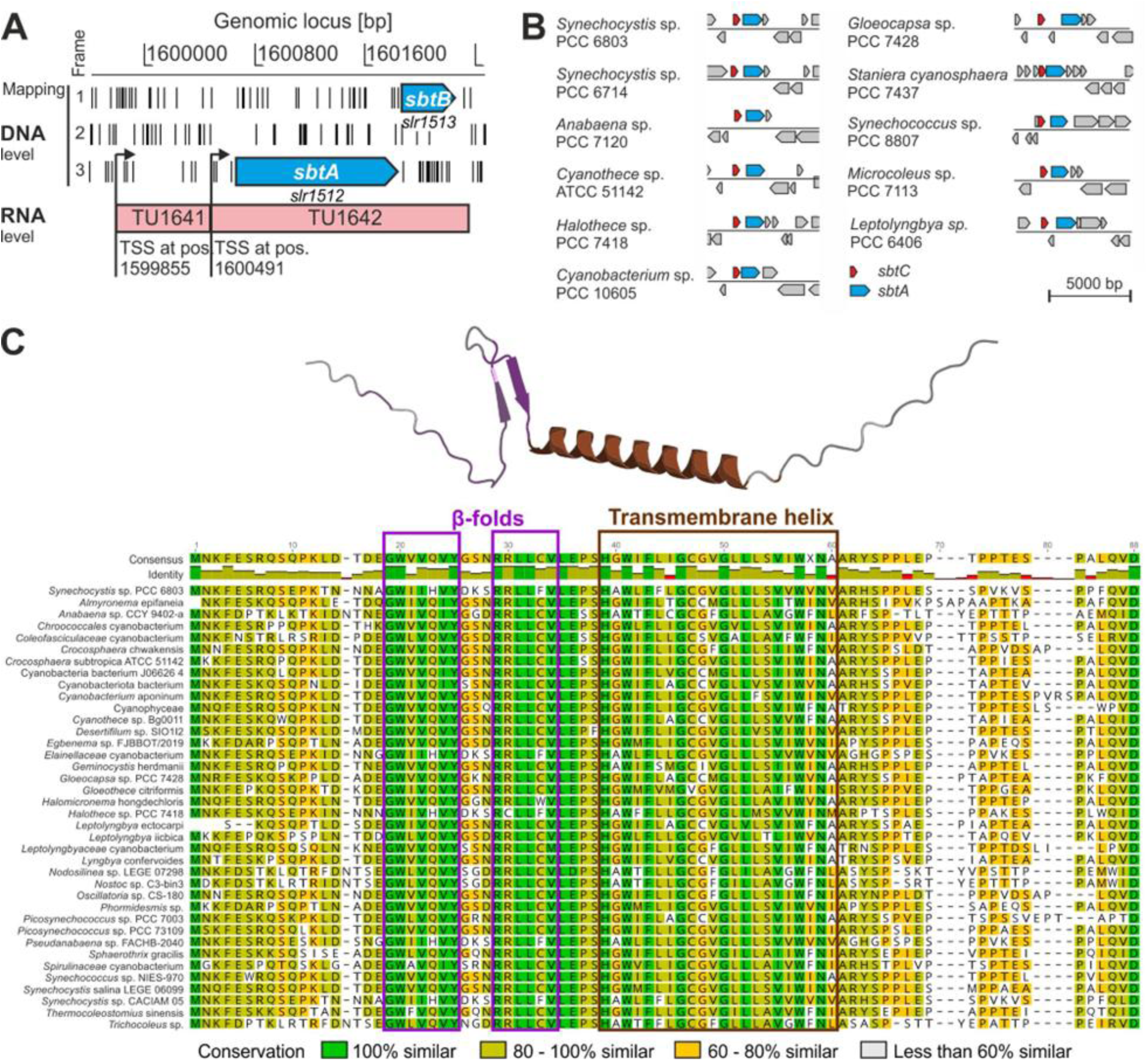
A non-annotated protein-coding gene upstream of *sbtA* in *Synechocystis* and various other cyanobacteria. **A:** Genomic region encoding the sodium-dependent bicarbonate transporter SbtA and the regulatory PII-like protein SbtB. The mapping of transcriptional start sites (TSS) and transcriptional units (TU) was extracted from a previous transcriptome analysis (Kopf *et al*. 2014), see also **Fig. S2**. The distribution of stop codons (shown by black bars) indicates that TU1641 harbors an open reading frame, which potentially encodes a protein of 80 amino acids. **B:** Corresponding *sbtC-sbtA* loci exist in in many recently published cyanobacterial genomes. In most genomes, both genes were found adjacent to each other, indicating their association. **C:** Multiple sequence alignment of SbtC homologs from 38 different cyanobacteria with different morphologies and lifestyles. The alignment shows a high level of conservation in the central region of the protein. Structure predictions using AlphaFold indicated that these regions form two β-sheets (purple) and a transmembrane helix (brown).

Interestingly, although several other bacteria outside the cyanobacterial phylum harbor *sbtA* genes, *sbtC* appears to be restricted to cyanobacteria, suggesting that it plays a specific role in cyanobacterial Ci assimilation. Moreover, it should be noted that not all cyanobacteria that harbor SbtAB also possess a SbtC homolog, which points toward a role that might be linked, but is not essential for SbtA function. For example, SbtC appears to be absent from *Synechococcus elongatus* and the picoplanktonic clades of α-cyanobacteria, which mainly include *Synechococcus* spp. and *Prochlorococcus* spp.. The latter cyanobacteria, however, underwent substantial genome reductions, resulting in the deletion of many, particularly regulatory proteins, from their genomes (Scanlan *et al*. 2009). Furthermore, they seem to harbor only distantly related SbtA2 proteins (Rae *et al*. 2011), which may function differently from the well-characterized SbtA proteins, then called SbtA1 (Rourke *et al*. 2025). Altogether, SbtC homologs could be identified in more than 140 cyanobacterial genomes, suggesting an important and conserved function potentially associated with the CCM.

### The sbtC locus is transcribed and stimulated by Ci limitation

Previous microarray analysis of *Synechocystis* cells shifted from high to low CO_2_ environment (Klähn *et al*. 2015) indicated enhanced transcription of the DNA region harboring *sbtC* under Ci limitation similar to *sbtAB* (**Fig. 2A**). In addition, former transcriptome analyses (Kopf *et al*. 2014) had indicated induction of transcription under Ci limitation and after transfer to high light (**Fig. S2**), which likely had triggered a higher Ci demand secondarily. Therefore, northern blots were performed using radiolabeled probes specific either for *sbtA* or *sbtC* to confirm that the genomic region upstream of *sbtAB* is indeed transcribed in a Ci-dependent manner (**Fig. 2B**). In presence of elevated CO_2_ (5% CO_2_ [v/v], high CO_2_, HC conditions) no transcripts were detected in *Synechocystis* wild type (WT). However, when cells were shifted to ambient air (0.04% CO_2_ [v/v], low CO_2_, LC conditions), indeed a signal that roughly corresponded to the estimated length of ∼600 nt for TU1641 was detected when using a probe specific for *sbtC*. The expression levels reached their maximum 12 h after the LC shift. The kinetics were comparable with those of the *sbtAB* operon, for which an abundant transcript of ∼2000 nt length was detected (**Fig. 2B**). The different sizes confirmed that *sbtC* is transcribed independently from *sbtA,* consistent with the two previously mapped transcriptional start sites at genomic positions 1,599,855 (*sbtC*, TU1641) and 1,600,491 (*sbtAB*, TU1642) (see **Fig. 1A, Fig. S2**). These findings unambiguously showed the inducibility of *sbtC* expression by LC conditions. Thus, the notion was supported that SbtC is somehow related to Ci assimilation or its regulation, as has been shown for SbtA and SbtB.

**Figure 2:**
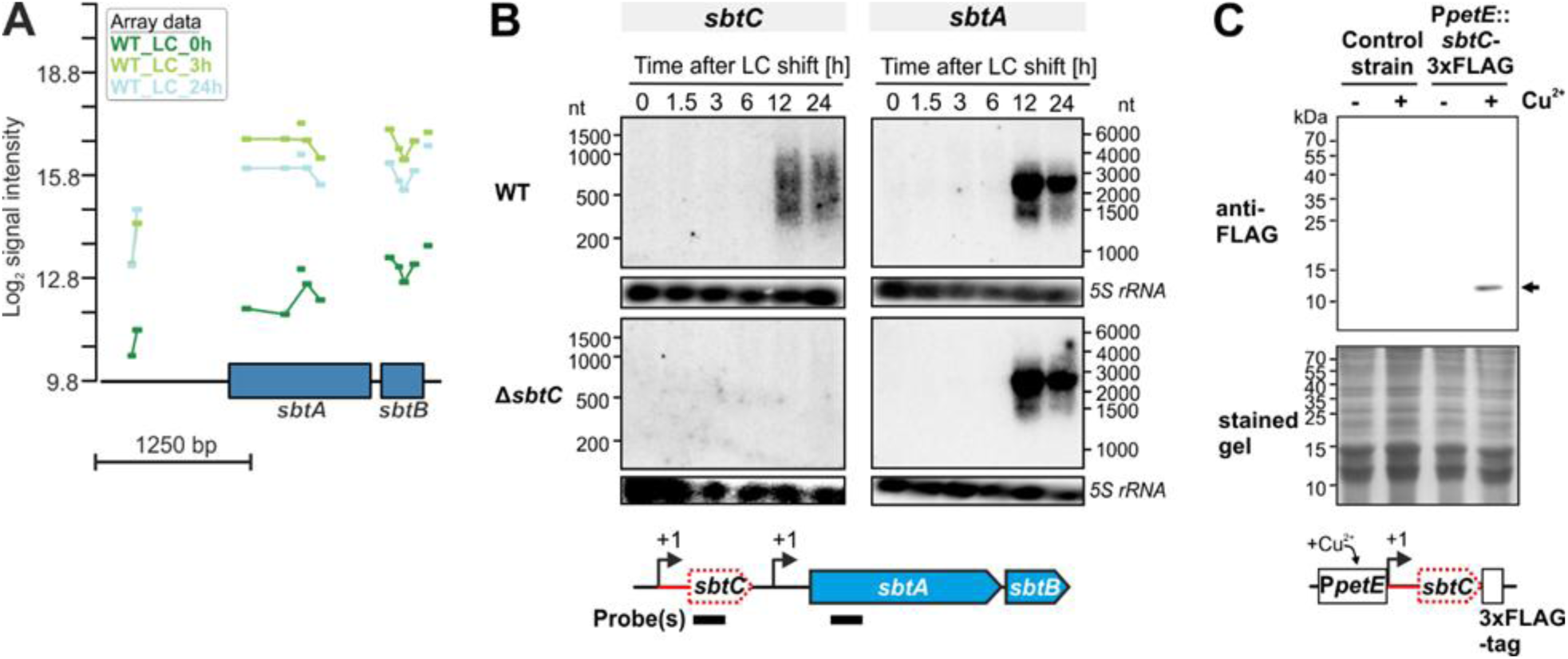
Verification of Ci-regulated *sbtC* expression in *Synechocystis*. **A:** Microarray data of *Synechocystis* cells, which were transferred from high CO_2_ (HC, 5% CO_2_ [v/v]) to low CO_2_ (LC, 0.04% CO_2_ [v/v]) and incubated for 3 and 24 h, respectively. Cells that were sampled before the CO_2_ downshift (0 h) served as a reference. Data are given as log_2_ signal intensities for probes covering the *sbtAB* genes and parts of the upstream region for which no gene was mapped. Probes are given as colored bars. The array data were extracted from a previous publication (Klähn *et al*. 2015). **B:** Northern blots confirming transcription of the region upstream of *sbtA*, which represents the yet non-annotated *sbtC* gene. The *sbtC* gene is clearly up-regulated in response to downshifts in Ci supply (from HC to LC) in WT and the transcript is independent from *sbtA*. Thus, a deletion of *sbtC* does not affect *sbtA* gene expression (compare signals for *sbtA* in WT and Δ*sbtC*). 5S rRNA was used as loading control. The lower panel illustrates the genomic organization and the sequence location that is covered by the probes used for the Northern blots (indicated by black bars). **C:** Western blot confirming the synthesis of a 3x-FLAG-tagged SbtC protein from this transcript. Transcription was driven by the Cu^2+^-inducible *petE* promoter, but translation initiation still relies on the 189 nt long native 5′-UTR upstream of the *sbtC* start codon (see also **Fig. S2**). Accordingly, the TU1641 represents an mRNA encoding a SbtC protein.

Next, we aimed at verifying that *sbtC* is indeed encoding a protein and does not represent a long non-coding RNA. To this end, we fused the gDNA covered by TU1641, i.e., the region between its transcriptional start site and the next one (TU1642 that covers *sbtA*) with a controllable promoter. In this case, we used the *petE* promoter, which responds to micromolar concentrations of copper ions. In this way, transcription of the genetic construct can be induced by the addition of Cu^2+^ but translation initiation still relies on the native sequences present within TU1641 (e.g., a putative Shine-Dalgarno sequence). To detect a potentially generated protein, the *sbtC* ORF was additionally fused to a 3xFLAG-tag encoding sequence (**Fig. 2C**). This construct was introduced into *Synechocystis* WT and respective cell lines were cultivated in the presence (+) or absence (-) of Cu^2+^. Indeed, by using an anti-FLAG antibody a protein that corresponds to the expected size was detected only when transcription was induced by Cu^2+^ (**Fig. 2C**). Consequently, the transcript must contain sequences required for translation initiation and hence, TU1641 indeed represents an mRNA encoding the SbtC protein.

### Functional analysis of SbtC in wild type

Due to its phylogenomic association with the *sbtAB* genes as well as its Ci-dependent regulation it was tempting to speculate that SbtC might represent a novel functional and/or regulatory component of the known Ci uptake systems, i.e. SbtA. To investigate its function, a knockout strain (Δ*sbtC*) was generated, in which no *sbtC* transcript could be detected (**Fig. 2B**). In contrast, the *sbtA* gene showed the same expression as in WT, confirming their independent transcription. Growth of the Δ*sbtC* mutant was compared to WT under HC as well as LC conditions. A drop dilution test on BG11 agar plates revealed no obvious growth deficit for the Δ*sbtC* strain regardless of the cultivation conditions (**Fig. 3A**). This finding is consistent with the observation of unchanged growth under LC of a Δ*sbtA* mutant (Shibata *et al*. 2002), because cyanobacterial Ci uptake is redundant due to the existence of multiple independent uptake systems (Burnap, Hagemann and Kaplan 2015; Hagemann, Song and Brouwer 2021).

**Figure 3:**
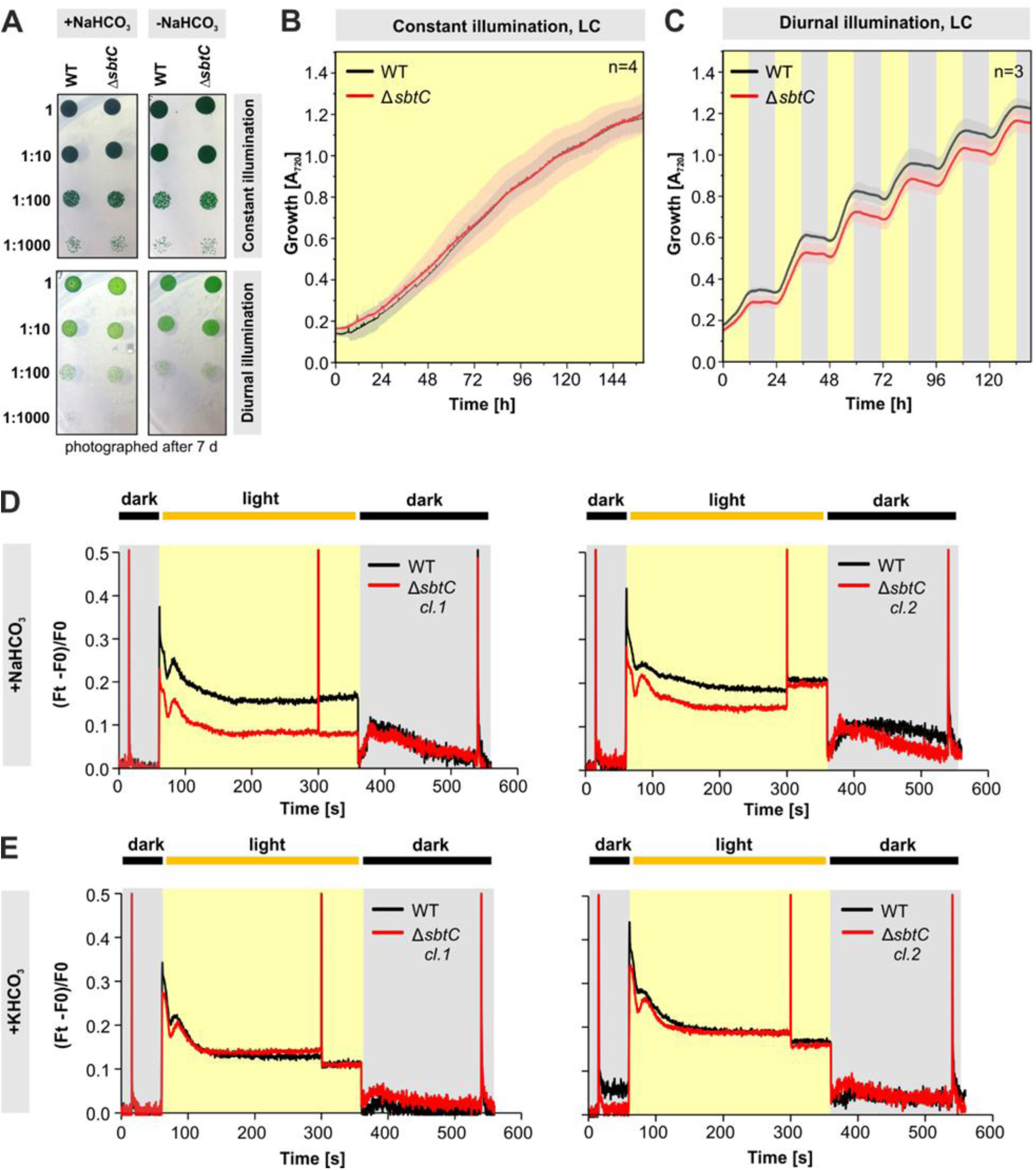
Phenotyping of a Δ*sbtC* mutant compared to WT. **A:** Photograph of a drop dilution assay. Both wild-type and Δ*sbtC* strains were cultivated on BG11 agar plates under four combinations of illumination (diurnal or constant) and supplementation (+/-NaHCO_3_). Photographs were taken after 7 days of cultivation. **B, C:** Growth performance of WT and Δ*sbtC* cells in Multi-Cultivator MC 1000. Cells were grown in BG11 medium under continuous (**B**) or diurnal (**C**) illumination (indicated by yellow and grey background) at 30°C and ambient CO_2_. Data are the mean ± standard deviation (displayed as pale area around the curves) of three biological replicates. **D, E:** Pulse Amplitude Modulation (PAM) fluorometry measurements using two different clones of Δ*sbtC*. Prior to the assay, cells were supplemented with 10 mM sodium (**D**) or potassium bicarbonate (**E**).

After obtaining these data, we analyzed the Δ*sbtC* mutant in more detail using a Multi-Cultivator MC 1000 (Photon Systems Instruments) that allows automatic optical density (OD) monitoring and hence, growth analysis at a higher resolution. In addition to continuous light conditions, we also monitored growth under a 12 h light/12 h dark regime. Furthermore, several Ci conditions were tested. Under continuous light, Δ*sbtC* again showed no difference in growth compared to the WT (**Fig. 3B**). Identical observations were made when cultivating WT and Δ*sbtC* under HC conditions (data not shown). Interestingly, when illuminated diurnally, the Δ*sbtC* mutant exhibited a slightly reduced growth relative to the WT (**Fig. 3C**).

Moreover, energy dissipation from the photosynthetic apparatus was affected in Δ*sbtC* (**Fig. 3D**). When shifting dark-adapted *Synechocystis* cells into the light, we observed a significantly lower chlorophyll a fluorescence in the Δ*sbtC* strain suggesting a greater capacity of converting light into chemical energy. This effect was linked to the presence of sodium bicarbonate, as no difference in chlorophyll a fluorescence could be observed in the presence of potassium bicarbonate (**Fig. 3E**). Again, this finding indicated a link to SbtA-mediated Na^+^-bicarbonate uptake, given that SbtA requires adequate amounts of sodium ions for bicarbonate symport. Altogether, these data suggest that the newly discovered SbtC protein plays an important role in Ci assimilation, where it not only cooperates with SbtA, but also with SbtB.

### Functional analysis of SbtC in a mutant strain expressing only sbtA for Ci uptake

The redundancy of several bicarbonate uptake systems made it difficult to clearly analyze how SbtC might specifically interact and regulate SbtA in cooperation with SbtB under different Ci levels. Therefore, we generated strains in the background of a previously generated Δ5 mutant (Xu *et al*. 2008), in which all five Ci transport systems of *Synechocystis* are mutated (**Fig. 4A**). A similar approach was recently reported to characterize BicA homologs among picocyanobacteria (Nieves-Morión *et al*. 2025). Due to the severe phenotypic effects of lacking these systems, the Δ5 strain can only be cultivated under HC conditions. However, a complementation strain Δ5::*sbtA* in which the mutated *sbtA* gene copy was replaced by the native *sbtA* gene via homologous recombination was able to grow at ambient air due to the sole activity of SbtA (**Fig. 4B**). This strain subsequently served as a platform to separately knock out either the *sbtB* or the *sbtC* genes by insertion of an erythromycin or gentamicin cartridge, respectively. The completely segregated strains Δ5::*stbAΔsbtB* and Δ5::*stbAΔsbtC* still grew at LC, indicating that neither protein is essential for SbtA-mediated bicarbonate uptake or its photosynthetic utilization (**Fig. 4B**).

**Figure 4.**
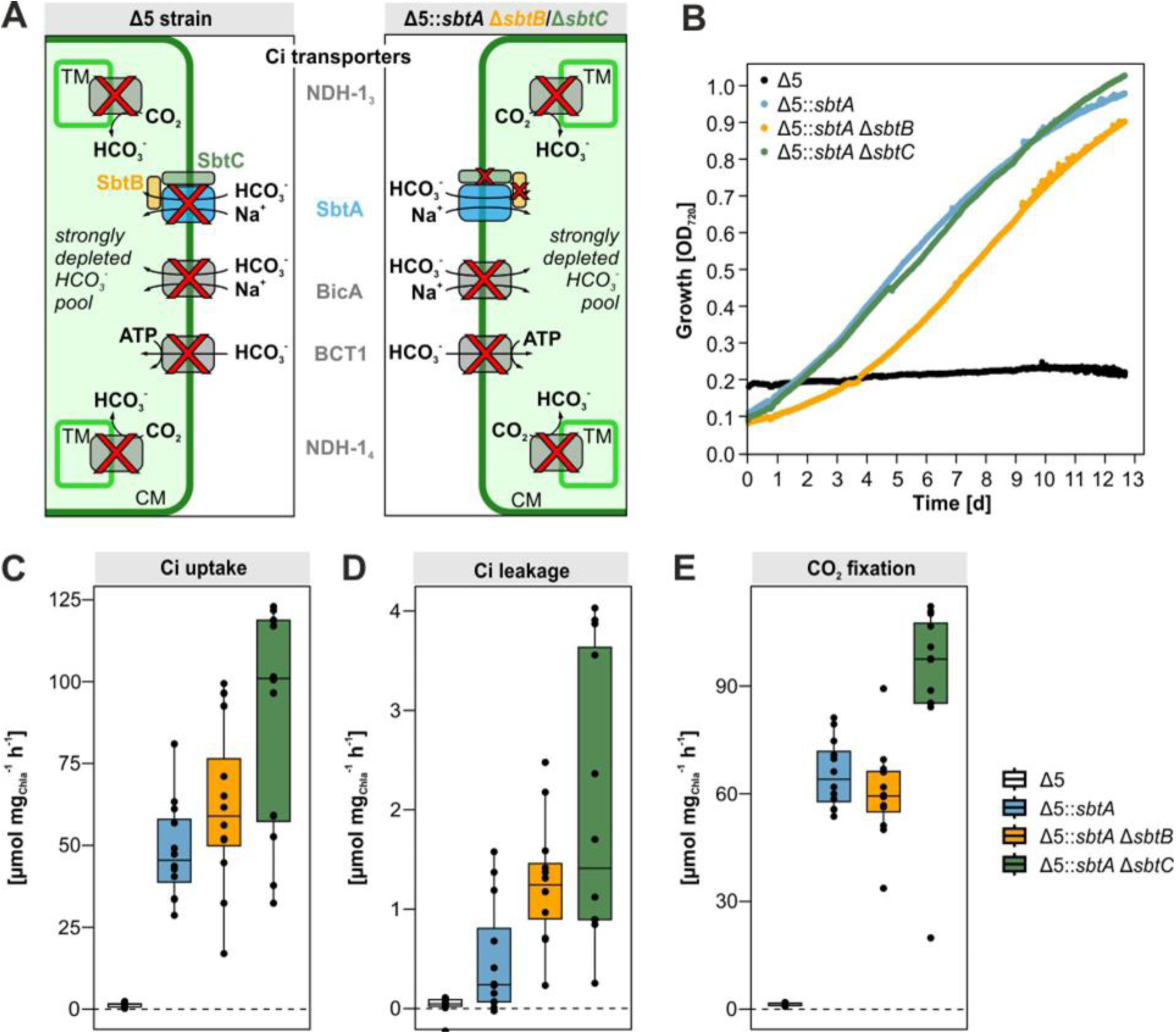
Inorganic carbon (Ci) uptake, leakage, and assimilation in different *Synechocystis* strains. **A:** Ci transport in the Δ5-derived strains. Δ5 lacks all Ci transporters. In Δ5::*sbtA*, the transporter SbtA is reintroduced; Δ5::*sbtAΔsbtB* and Δ*sbtC* lack either of the proteins that potentially regulate bicarbonate transport via SbtA. **B:** Growth curves of the Δ5-derived strains at ambient air (0.04% CO_2_ [v/v]) and constant illumination. Δ5 is not vital under ambient air conditions. The reintroduction of SbtA rescues this lethal phenotype. 50 µmol photons m^-2^ s^-1^. **C:** The initial mutant Δ5 did not take up significant amounts of Ci during the experimental time frame. Strains Δ5::*sbtA* and Δ5::*sbtA*Δ*sbtB* did not show a significant difference in Ci uptake (Welch-Two Sample Test, p = 0.309). There is neither statistical significance between Δ5::*sbtA*Δ*sbtB* and Δ5::*sbtA*Δ*sbtC* (p = 0.058). Strain Δ5::*sbtA*Δ*sbtC* showed significantly higher Ci uptake than Δ5::*sbtA* (p = 0.007). The Δ5 strain did not show leakage because it did not take up Ci in the first place. Δ5::*sbtA* showed little leakage, while Δ5::*sbtA*Δ*sbtB* leaked significantly more (p = 0.004), corresponding to (Haffner *et al*. 2023). Interestingly, Δ5::*sbtA*Δ*sbtC* exhibited the same phenotype (comparing AC p = 0.004, comparing BC p = 0.111). While Δ5 only showed little Ci assimilation (acid-stable ^14^C), similar CO_2_ fixation rates were determined for Δ5::*sbtA* and Δ5::*sbtA*Δ*sbtB* (p = 0.249). On the other hand, Δ5::*sbtA*Δ*sbtC* showed significantly higher assimilation rates than Δ5::*sbtA* (comparing AC p = 0.004, comparing BC p = 0.001); n=12.

The three generated strains, expressing either *sbtA* together with both native *sbtB* and *sbtC* loci (Δ5::*sbtA*), *sbtA* with only *sbtC* (Δ5::*sbtAΔsbtB*), or *sbtA* with only *sbtB* (Δ5::*sbtAΔsbtC*), permitted a detailed analysis of SbtA-related functions in the presence of both or just one of the likely regulatory proteins. Incubation of the strains with ^14^C-labelled bicarbonate allowed measurement of three parameters directly related to SbtA function. The uptake of ^14^C-labelled bicarbonate reflects the overall SbtA transport activity, while the incorporation of label into acid-stable metabolites displays overall photosynthetic carbon fixation. Both Ci uptake and CO_2_ fixation were negligible in the Δ5 mutant control. At the same time, no differences were detected in the two parameters between the strains Δ5::*sbtA* and Δ5::*sbtAΔsbtB* (**Fig. 4C**). However, in Δ5::*sbtAΔsbtC*, significantly higher Ci uptake was observed, pointing towards a negative impact of SbtC on SbtA-dependent bicarbonate uptake. Additionally, the strain Δ5::*sbtAΔsbtC* showed significantly higher CO_2_ fixation (**Fig. 4**), which corresponded to higher amounts of RbcL protein per chlorophyll a (**Fig. S3**). Finally, Haffner *et al*. (Haffner *et al*. 2023) demonstrated that SbtA acts as a channel that leaks bicarbonate in darkness, while SbtB functions as a plug to minimize this leakage. This finding was confirmed by our experiments, where we observed significantly higher bicarbonate release in the strain Δ5::*sbtAΔsbtB* than in Δ5::*sbtA*. Furthermore, the results indicated that the SbtB plug depends on the interaction with SbtC to fulfil this task, because a similarly increased leakage was observed in strain Δ5::*sbtAΔsbtC* in which *sbtB* remained intact (**Fig. 4C**).

### SbtC is located in the cytoplasmic membrane and interacts with SbtA and B

Multiple sequence alignment of SbtC homologs from several different cyanobacteria revealed a highly conserved central region within the protein. Structural predictions indicate that this region may form a transmembrane helix and two β-sheets with cytoplasmic localization. At the same time, the less conserved N-terminal and C-terminal parts seem to be mostly unstructured (**Fig. 1C**). To verify this proposed membrane-associated localization and function, the strain Δ*sbtC*+pVZ321-P*petE*::*sbtC*::3xFLAG expressing FLAG-tagged SbtC was analyzed by Blue Native (BN) PAGE. Distinct membrane complexes, including trimeric photosystem I and dimeric photosystem II, were resolved (**Fig. 5**). Immunoblotting with SbtA and SbtB-specific antibodies revealed that both proteins co-migrated within the same high-molecular mass complexes, verifying their direct interaction. Notably, the small FLAG-tagged SbtC was detected at the same two positions as the SbtA-SbtB protein complexes (**Fig. 5**). These findings point to the existence of a ternary SbtABC complex in the cytoplasmic membrane of *Synechocystis*. The lower bands seem to correspond to the expected size of the ternary SbtABC complex of approximately 180 kDa (SbtA trimer 120 kDa, SbtB trimer 36 kDa, SbtC-Flag 12 kDa). This assumption is supported by its migration similar to the Cytb6f complex, which is approximately 170 kDa (**Fig. 5**). The upper band with the predominant SbtA signal migrated just below the PSII monomer that has a size of approximately 230 kDa. Hence, the upper bands may represent a supercomplex with more SbtA subunits or other interacting proteins together with SbtB and SbtC.

**Figure 5:**
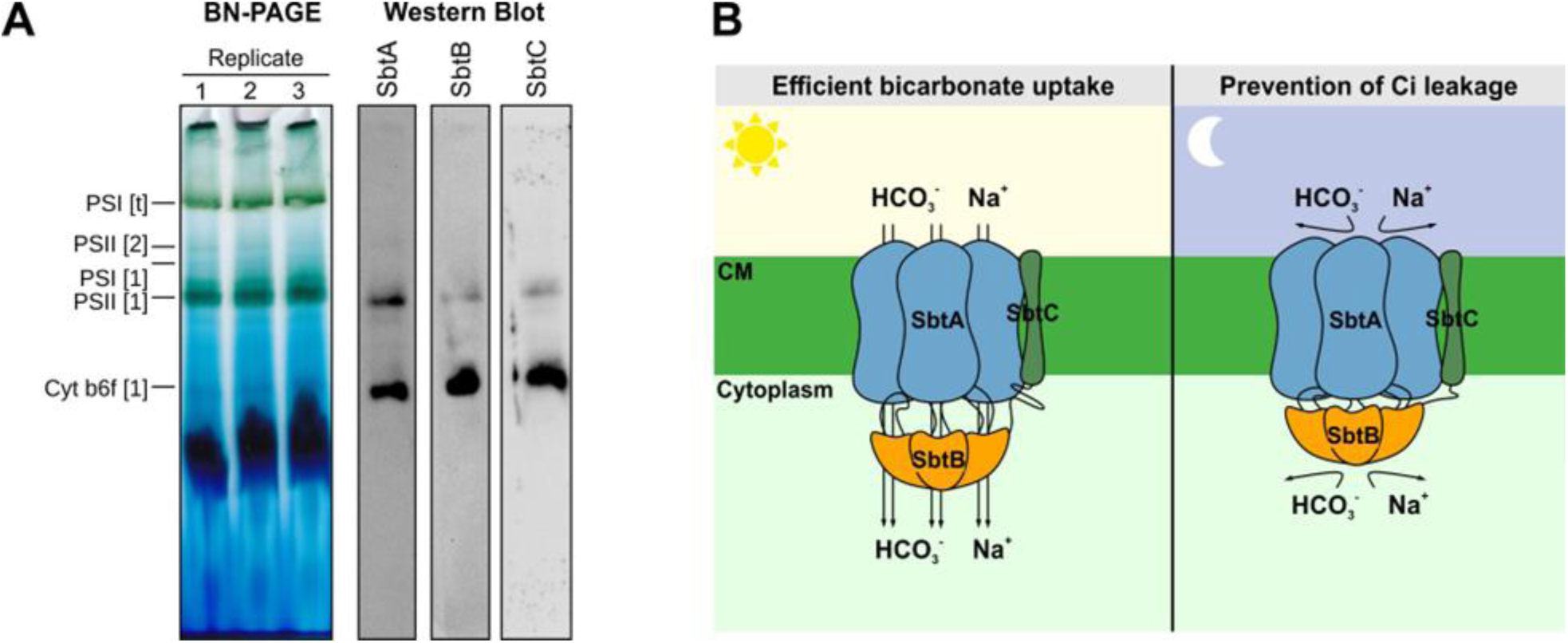
Existence of a ternary complex of SbtC with SbtA and SbtB. **A:** Left panel lanes: Picture of the separation of enriched membranes from *Synechocystis* in BN gels. Annotation of the colored bands in the BN gel according to (Casella *et al*. 2017). Right panel lanes: Blotted BN gels were probed with specific antibodies for SbtA, SbtB, or the FLAG-tag fused to SbtC. Two high-molecular-mass complexes at approximately 180 kDa and 220 kDa were recognized by the three antibodies. **B:** Proposed interaction mechanism of SbtA, B, and C. During the day, SbtB and SbtC stabilize SbtA, thereby enabling efficient bicarbonate uptake. At night, the combination of SbtB and SbtC prevents bicarbonate leakage out of the cell, with SbtB acting as the main plug and SbtC stabilizing the binding of SbtB to SbtA. Simultaneously, bicarbonate import is prevented. SbtC assists specifically with day-night transitions of SbtA activation and inactivation. Information about the trimeric structures of SbtA and B has been adopted from structural studies (Liu *et al*. 2021).

## Discussion

In our study, we show that in *Synechocystis* and many other cyanobacteria, a novel, previously not annotated protein-coding gene exists upstream of the well-conserved *sbtAB* operon. This is surprising as the SbtA/B pair received a lot of attention during the last years due to its crucial function in Ci assimilation. However, this finding is not uncommon, as many open reading frames (ORFs) encoding small proteins (≤80 or ≤100 amino acids) have systematically been overlooked in bacteria (e.g., (Storz, Wolf and Ramamurthi 2014; Baumgartner *et al*. 2016)). In recent years, with the emerging research on small proteins, their crucial role in many important cellular processes has been increasingly recognized, including those of cyanobacteria (Brandenburg and Klähn 2020; Burton, Zeinert and Storz 2024; Kraus and Hess 2025). Due to their small size, a direct enzymatic function is unlikely, but they may instead represent i) structural subunits of larger complexes; ii) regulatory factors in gene expression, or iii) regulatory subunits of proteins/enzymes connecting them with others (Hobbs *et al*. 2011; Gray, Storz and Papenfort 2022). In the case of the cyanobacterium *Synechocystis*, small proteins have been assigned as regulators of gene expression (e.g., PipX (Espinosa *et al*. 2006)), enzyme activity regulators (e.g., PirA (Bolay *et al*. 2021), PirC (Orthwein *et al*. 2021), NirP1 (Kraus *et al*. 2024), AcnSP (de Alvarenga, Hess and Hagemann 2020)), and/or structural components of larger protein complexes (AtpΘ (Song *et al*. 2022), NdhP (Schwarz *et al*. 2013; Zhang *et al*. 2014), RbcX (Saschenbrecker *et al*. 2007)).

In our study, we demonstrate that the small protein SbtC encoded upstream of the *sbtAB* operon is co-regulated with other CCM genes, i.e., it is upregulated in response to Ci limitation. However, data from previous transcriptome analyses indicated that *sbtC* expression appears not to be regulated by the repressor NdhR, as, in contrast to *sbtA*, *sbtC* was not de-repressed in Δ*ndhR* mutants under high CO_2_ (Klähn *et al*. 2015) (corresponding data provided in **Supplementary Fig. S4A**). The latter further confirmed that *sbtC* is transcribed independently from *sbtAB*. Inspection of other available transcriptome datasets suggested that *sbtC* might be regulated by RbcR, the other LysR-type transcriptional regulator involved in the transcriptional regulation of CCM genes (Bolay *et al*. 2022) (corresponding data provided in **Supplementary Fig. S4B**). In a partial *rbcR* knockout strain, the *sbtC* transcript levels were clearly decreased in comparison to WT expression, similar to *sbtAB*. Apparently, full expression of both *sbtAB* and *sbtC* requires sufficient amounts of RbcR. Moreover, LC-induced expression of both *sbtC* and *sbtA* is affected in a Δ*sbtB* strain, with both genes showing reduced transcript abundance (Mantovani *et al*. 2022) (corresponding data provided in **Supplementary Fig. S4C**). This could point towards the involvement of the PII-like protein in the positive regulation of *sbtC* and *sbtA* via an as-yet-unidentified mechanism.

Consistently, *sbtC* deletion affects the growth of *Synechocystis* under diurnal conditions similar to the previously reported effects of the *sbtB* mutant (Selim *et al*. 2021), suggesting that these proteins may jointly contribute to the acclimation of cellular processes to naturally fluctuating light regimes, such as day-night cycles. In a specific mutant background in which the growth on Ci depends solely on the SbtA activity, we further demonstrate that SbtC significantly influences SbtA-dependent effects such as bicarbonate uptake and bicarbonate leakage through SbtA during darkness. Bicarbonate leakage in darkness has also been observed before in the absence of SbtB in the WT background, which led to the formulation of the “plug hypothesis” for SbtB and related PII-like proteins (Haffner *et al*. 2023). Finally, SbtC was detected as a part of membrane-associated complexes together with SbtA and SbtB (see **Fig. 5**), consistent with a potential direct interaction of the proteins. Collectively, these findings strongly indicate that the 80-amino-acid protein SbtC is closely associated with SbtA and/or SbtB and may play an important role in modulating SbtA activity and stability, particularly under fluctuating light and Ci conditions.

The limited length of the protein, and hence the absence of any conspicuous domains, make predictions on its specific function challenging. Structure predictions show that the well conserved protein possesses a central transmembrane helix, consistent with its likely membrane localization. Its presence in the prepared membrane complexes further corroborated this prediction. In addition to the central helix, the N-terminal part of SbtC is predicted to form two highly conserved two β-sheets, which might be relevant for specific protein-protein interactions. One likely candidate is SbtB, which lacks an own transmembrane domain and was found to be associated with the membrane or as a soluble protein in *Synechocystis* depending on the Ci conditions (Selim *et al*. 2018). Impaired association of SbtB with the SbtA bicarbonate transporter likely explains the overlapping phenotypic alterations in the *sbtB* and *sbtC* mutants, i.e. the inability to prevent bicarbonate leakage and the slower growth at diurnal conditions.

While the prevention of bicarbonate leakage in the dark is directly linked to the function of SbtA as predicted previously (Haffner *et al*. 2023) and confirmed here in our mutant system with only SbtA as bicarbonate transporting pore, the decreased diurnal growth is rather linked to the role of SbtB in regulating GlgB activity for correct glycogen metabolism (Selim *et al*. 2021). Nevertheless, we cannot exclude that darkness-induced loss of bicarbonate might also contribute to the weaker growth of the knock-out strain under diurnal conditions. Interestingly, SbtC seems to positively influence the bicarbonate uptake via SbtA directly, because its absence stimulated the transport activity, whereas SbtB is rather not involved in this function. The slight increase of bicarbonate fixation seems to be rather related to the high RubisCO content (see **Suppl. Fig. S3**) in the strain with mutated *sbtC* genes, which might represent an indirect effect.

A direct regulatory role of SbtC on *sbtAB* expression seems unlikely, as *sbtAB* transcript accumulation is similar in the *sbtC* knock-out strain and in the WT. However, we cannot rule out that SbtC might also interact with transcription factors contributing to the Ci-regulated expression of *sbtAB* or other CCM-related genes. For example, it has been reported that the cAMP-dependent transcriptional factor SyCRP can be membrane-associated or not, which clearly depends on the Ci conditions for growth (Bantu *et al*. 2022). Similar observations have been reported for the membrane association of SbtB (Selim *et al*., 2018). Hence, it is possible that the SbtC protein could be involved in the formation of a Ci-related complex with SbtA sequestering SbtB and/or SyCRP. The occurrence of the second larger SbtABC complex exceeding the theoretical size of kDa of a solely SbtA[3]B[3]C_FLAG_[1-3] supercomplex might give a hint that the SbtABC complex could represent a hub where also other CCM-related proteins, possibly SyCRP or others might be anchored.

Clearly, additional experiments are needed to fully elucidate the precise role of SbtC and its interaction with SbtA and possible other targets. Isolation and structural characterization of the ternary SbtABC complex would represent a significant breakthrough, although such analyses are beyond the scope of the present paper.

## Material and Methods

### Strains and cultivation

A glucose-tolerant strain of *Synechocystis* sp. PCC 6803 (substrain Kazusa, obtained from N. Murata, National Institute for Basic Biology, Okazaki, Japan) was used as the WT. All cyanobacterial strains were cultivated at 30°C in BG11 medium (Rippka *et al*. 1979) that was slightly modified as follows: it was buffered with 20 mM TES–KOH to pH 8.0 and did not contain Na_2_CO_3_. For cultivation on plates, BG11 minimal medium was solidified with Bacto^TM^ agar (15 g l^-1^, Fisher Scientific) and supplemented with 3 g l^-1^ Na_2_S_2_O_3_ (Thiel, Bramble and Rogers 1989). Growth was conducted in a plate incubator set to 1% CO_2_, 30 °C, and 75% humidity, under continuous white light illumination of 50 μmol photons m^−2^ s^−1^. For liquid cultures, cells were grown in Erlenmeyer flasks shaken at 150 rpm under conditions otherwise identical to plate cultures.

Mutant strains were selected and maintained in the presence of respective antibiotics with the following concentrations: 50 µg/ml kanamycin, 2.5 µg/ml gentamicin, 40 µg/ml hygromycin, 10 µg/ml streptomycin, 10 µg/ml chloramphenicol or 50 µg/ml erythromycin.

To characterize growth performance of the obtained mutant strains compared to the WT, cultivation was performed in a Multi-Cultivator MC 1000-OD (Photon Systems Instruments, Czech Republic) using BG11 medium without antibiotics. OD_720_ was measured in 10-minute intervals. Growth was monitored at different CO_2_ or light conditions. To establish a low CO_2_ environment (LC conditions) cells were bubbled with ambient air containing 0.04% CO_2_, whereas at high CO_2_ (HC conditions), the air stream was enriched with 2% CO_2_ (v/v). Additionally, the influence of illumination modes on growth was assessed by cultivating under either constant illumination with 50 μmol photons m^−2^ s^−1^ or under a diurnal light regime (12h light/12h dark).

### Mutant generation

The oligonucleotides used to generate the mutant strains are given in **Supplementary Table S1**, all strains used in this study are given in **Supplementary Table S2**. The Δ*sbtC* single mutant was obtained by homologous recombination, i.e. partial replacement of the corresponding open reading frame by a gentamicin resistance cassette (Gm^R^). For this, flanking regions present up-and downstream of the previously annotated TU1641 (Kopf *et al*. 2014) were amplified from *Synechocystis* gDNA using primer pairs 1641_up_fw/_rev and 1641_down_fw/_rev. Sites for the restriction endonucleases NdeI and NotI were added to the primers and allowed ligation with a similarly obtained PCR product harboring Gm^R^ that was amplified using primers Gm_fw_NotI/Gm_rev_oop_NdeI. Via the latter PCR an oop terminator sequence was also introduced downstream of Gm^R^. After digestion of the PCR products with NdeI and NotI, the fragments were ligated using T4 ligase. All enzymes were obtained from Thermo Fisher Scientific and used according to the manufacturer’s protocols. The final construct was amplified from the obtained ligation product using primers 1641_seq_fw/_rev and introduced into pJET1.2 using the CloneJET PCR Cloning Kit (Thermo Fisher Scientific). The corresponding plasmid was used to transform *Synechocystis* WT and transformants were selected on 1 µg/ml gentamicin.

The CCM-deficient mutant Δ5 was described previously (Xu *et al*. 2008). This mutant cannot survive under ambient air conditions and, therefore, was cultivated at elevated CO_2_ conditions (2% v/v). This mutant was restored for SbtABC function via transformation with DNA harboring the genetic locus of *sbtABC* and selecting for growth at ambient air conditions resulting in the mutant Δ5::*sbtA*. The corresponding DNA was obtained by PCR using gDNA of the WT as template and primers Psbt_fw and Psbt_rv. In the mutant Δ5::*sbtA*, additional knockouts for *sbtB* and *sbtC* were established. Knockout of *sbtB* resulting in strain Δ5::*sbtA*Δ*sbtB* was achieved by the transformation of Δ5::*sbtA* with a PCR fragment amplified from gDNA of a single Δ*sbtB* mutant (Selim *et al*. 2018) using primers Psbt_fw and Psbt_rv. Transformants were selected for growth at ambient air and erythromycin resistance. For comparison reasons (and as Gm resistance is already present in Δ5), an erythromycin resistance cassette was also used to achieve a separate knockout of *sbtC* in Δ5::*sbtA.* The corresponding construct was obtained by fusion PCR using the primers P1-P6_sbtC^ery^. PCR genotyping showed fully segregated mutant strains (**Fig. S1**).

The constructs for Cu^2+^-inducible expression of FLAG-tagged SbtC variants were generated by gene synthesis (Integrated DNA technologies). Briefly, the genomic region representing the *sbtC* ORF and its 5′-UTR were fused to the promoter sequence of the *petE* gene from *Synechocystis* upstream and an *oop* terminator downstream of *sbtC*. The sequence of the final construct including descriptions is given in the Supplementary data. After gene synthesis the final constructs were re-amplified using primers PpetE-XhoI_fw and Toop-HindIII_rev, thereby adding XhoI and HindIII restriction endonuclease sites. The PCR product was cloned into pJET1.2 using the CloneJET PCR Cloning Kit (Thermo Fisher Scientific) from which it was subsequently cutted with XhoI and HindIII and introduced into the self-replicating vector pVZ321 (Zinchenko *et al*. 1999). The generated plasmid pVZ321-P*petE*::*sbtC*_3xFLAG or an empty pVZ321 plasmid were introduced via conjugation into *Synechocystis* WT and Δ*sbtC* resulting in strains WT+pVZ321, WT+pVZ321-P*petE*::*sbtC*::3xFLAG, Δ*sbtC*+pVZ321 and Δ*sbtC*+pVZ321-P*petE*::*sbtC*::3xFLAG.

### Physiological characterization of mutant strains

For PAM fluorescence measurements, a Walz DUAL-PAM 100 (Walz, Germany) with (DUAL-DR) and NADPH (DUAL-(E/D) NADPH) units was used and the experiments were conducted as described previously (Holland *et al*. 2016). The quantification of ^14^C-labeled inorganic carbon leakage was conducted as described previously (Haffner *et al*. 2023). All strains were pre-cultivated with CO_2_-enriched air (2%) and shifted overnight to ambient air before the experiment.

### Sequence analyses

Putative SbtC homologs were searched for using TBLASTN (Gertz *et al*. 2006). After manual inspection, encoded protein sequences were collected and sequence alignments were generated with ClustalW (Larkin *et al*. 2007).

### Blue-native gel electrophoresis

Membrane protein extracts were obtained from the strain Δ*sbtC*+pVZ321-P*petE*::*sbtC*::3xFLAG. After pre-cultivation in CO_2_-enriched air, the culture was shifted to ambient air for 20-24 h to induce *sbtAB* and *sbtC* expression. The cells were lysed in 25 mM MES/NaOH buffer (pH 6.5) containing 4 mM MgCl_2_, 4 mM CaCl_2_ and 25% (w/v) glycerol using a French Press Cell Disrupter (Thermo Electron Corporation, 6.9 MPa). The centrifuged (30 min, 14000 × g, 5°C) pellet was gently re-suspended in lysis buffer and stored at -80°C for analysis. For BN gels, cell extract equivalent to 8 µg chlorophyll a was suspended in buffer containing 750 mM aminocaproic acid, 50 mM BisTris/HCl (pH 7) and 0.5 mM Na_2_EDTA and solubilized using 1% dodecyl-β-D-maltoside for 30 min on ice. After centrifugation (10 min, 16000 × g, 5°C), gradient gels 4-16% (SERVAGel^TM^N) were run according to the manufacturer’s protocol. The voltage was successively increased by 25 V every 30 min from 50 V to 200 V. The blue cathode buffer was changed to clear buffer at approximately half the running distance.

### Northern and Western blotting

RNA extraction from *Synechocystis* was done as described previously (Hein *et al*. 2013). For Northern hybridization, 3 µg of total RNA was separated on denaturing agarose gels and transferred to Hybond-N+ membranes (Amersham, Germany) through capillary blotting with 20x SSC buffer (3 M NaCl, 0.3 M sodium citrate, pH 7.0). The membranes were hybridized with [α-^32P^]-UTP incorporated single-stranded RNA probes generated through *in vitro* transcription as previously described (Steglich *et al*. 2008). PCR fragments amplified from *Synechocystis* gDNA using primers pr_sbtA_fw/rev and pr_1641_fw/rev were used as templates for *in vitro* transcription of probes against *sbtA* and *sbtC*. After hybridization, the signals were detected using a Personal Molecular Imager system (Pharos FX, BIO-RAD, Germany) and analyzed using Quantity One software (BIO-RAD, Germany).

For the general detection of specific proteins in *Synechocystis*, total proteins were extracted from centrifuged cells using a Precellys homogenizer (Bertin Technologies). SDS-PAGE was performed in 12% polyacrylamide gels. Blotting on nitrocellulose membranes was done using an OWL Peqlab HEP-1 Semi-Dry Blotter (5 V, 1 h). For staining with ponceau red, the membrane was incubated for 5 min in a Ponceau solution and washed with dH_2_O for 1 min. A commercial antibody (Agrisera AS13 2657) was used for immunodetection of SbtA, and a custom-made antibody for SbtB (Selim *et al*. 2018). SbtC was detected by immunodetection of the FLAG-tag by ANTI-FLAG^®^M2 peroxidase antibody (Sigma-Aldrich). For blotting of BN gels a PVDF membrane was used, and proteins were transferred via vertical wet blot (120 V, 2 h).

## Acknowledgments

The UFZ is supported by the European Regional Development Funds (EFRE, Europe funds Saxony) and the Helmholtz Association. The project was funded by grants of the German Research Foundation (DFG) (grants HA2002/27-1 to MH; grant KL 3114/10-1 to SK and grant HE 2544/22-1 to WRH). The PhD thesis of PW was supported by the German Academic Scholarship Foundation (‘Studienstiftung des deutschen Volkes’). We thank Saskia Kürten for technical assistance at the University of Rostock. We also thank Marcus Ziemann at Freiburg University for the help with the RNAplot algorithm.

## Author contributions

S.K. designed the research; M.H. and S.K. supervised the project; P.W., C.P., N.M. and S.K. performed research; W.R.H. and R.B. contributed computational data or new analytic tools; S.K. and M.H. analyzed data; P.W., C.P., M.H. and S.K. wrote the paper. All authors approved the final version.

## References

de Alvarenga LV, Hess WR, Hagemann M. AcnSP - A novel small protein regulator of aconitase activity in the cyanobacterium *Synechocystis* sp. PCC 6803. Front Microbiol 2020;11:1445.

Bantu L, Chauhan S, Srikumar A et al. A membrane-bound cAMP receptor protein, SyCRP1 mediates inorganic carbon response in *Synechocystis* sp. PCC 6803. Biochim Biophys Acta Gene Regul Mech 2022;1865:194803.

Battchikova N, Eisenhut M, Aro E-M. Cyanobacterial NDH-1 complexes: novel insights and remaining puzzles. Biochim Biophys Acta 2011;1807:935–44.

Baumgartner D, Kopf M, Klähn S et al. Small proteins in cyanobacteria provide a paradigm for the functional analysis of the bacterial micro-proteome. BMC Microbiol 2016;16:285.

Bolay P, Rozbeh R, Muro-Pastor MI et al. The novel PII-interacting protein PirA controls flux into the cyanobacterial ornithine-ammonia cycle. mBio 2021;12, DOI: 10.1128/mBio.00229-21.

Bolay P, Schlüter S, Grimm S et al. The transcriptional regulator RbcR controls ribulose-1,5-bisphosphate carboxylase/oxygenase (RuBisCO) genes in the cyanobacterium *Synechocystis* sp. PCC 6803. New Phytol 2022;235:432–45.

Bowes G, Ogren WL, Hageman RH. Phosphoglycolate production catalyzed by ribulose diphosphate carboxylase. Biochem Biophys Res Commun 1971;45:716–22.

Brandenburg F, Klähn S. Small but Smart: On the Diverse Role of Small Proteins in the Regulation of Cyanobacterial Metabolism. Life (Basel) 2020;10, DOI: 10.3390/life10120322.

Burnap RL, Hagemann M, Kaplan A. Regulation of CO_2_ Concentrating Mechanism in Cyanobacteria. Life (Basel) 2015;5:348–71.

Burton AT, Zeinert R, Storz G. Large Roles of Small Proteins. Annu Rev Microbiol 2024;78:1–22.

Casella S, Huang F, Mason D et al. Dissecting the Native Architecture and Dynamics of Cyanobacterial Photosynthetic Machinery. Mol Plant 2017;10:1434–48.

Daley SME, Kappell AD, Carrick MJ et al. Regulation of the cyanobacterial CO_2_-concentrating mechanism involves internal sensing of NADP^+^ and α-ketogutarate levels by transcription factor CcmR. PLoS One 2012;7:e41286.

Du J, Förster B, Rourke L et al. Characterisation of cyanobacterial bicarbonate transporters in *E. coli* shows that SbtA homologs are functional in this heterologous expression system. PLoS One 2014;9:e115905.

Ellis RJ. Biochemistry: Tackling unintelligent design. Nature 2010;463:164–5.

Espinosa J, Forchhammer K, Burillo S et al. Interaction network in cyanobacterial nitrogen regulation: PipX, a protein that interacts in a 2-oxoglutarate dependent manner with P_II_ and NtcA. Mol Microbiol 2006;61:457–69.

Fang S, Huang X, Zhang X et al. Molecular mechanism underlying transport and allosteric inhibition of bicarbonate transporter SbtA. Proc Natl Acad Sci U S A 2021;118:e2101632118.

Figge RM, Cassier-Chauvat C, Chauvat F et al. Characterization and analysis of an NAD(P)H dehydrogenase transcriptional regulator critical for the survival of cyanobacteria facing inorganic carbon starvation and osmotic stress. Mol Microbiol 2001;39:455–68.

Flügel F, Timm S, Arrivault S et al. The photorespiratory metabolite 2-phosphoglycolate regulates photosynthesis and starch accumulation in *Arabidopsis*. Plant Cell 2017;29:2537–51.

Forchhammer K. Global carbon/nitrogen control by P_II_ signal transduction in cyanobacteria: from signals to targets. FEMS Microbiol Rev 2004;28:319–33.

Gertz EM, Yu Y-K, Agarwala R et al. Composition-based statistics and translated nucleotide searches: Improving the TBLASTN module of BLAST. BMC Biology 2006;4:41.

Gray T, Storz G, Papenfort K. Small Proteins; Big Questions. J Bacteriol 2022;204:e0034121.

Haffner M, Hou W-T, Mantovani O et al. PII signal transduction superfamily acts as a valve plug to control bicarbonate and ammonia homeostasis among different bacterial phyla. 2023:2023.08.10.552651.

Hagemann M, Fernie AR, Espie GS et al. Evolution of the biochemistry of the photorespiratory C2 cycle. Plant Biol (Stuttg) 2013;15:639–47.

Hagemann M, Song S, Brouwer E-M. Inorganic Carbon Assimilation in Cyanobacteria: Mechanisms, Regulation, and Engineering. *Cyanobacteria Biotechnology*. John Wiley & Sons, Ltd, 2021, 1–31.

Hammer A, Hodgson DRW, Cann MJ. Regulation of prokaryotic adenylyl cyclases by CO_2_. Biochem J 2006;396:215–8.

Hein S, Scholz I, Voß B et al. Adaptation and modification of three CRISPR loci in two closely related cyanobacteria. RNA Biol 2013;10:852–64.

Hobbs EC, Fontaine F, Yin X et al. An expanding universe of small proteins. Curr Opin Microbiol 2011;14:167–73.

Holland SC, Artier J, Miller NT et al. Impacts of genetically engineered alterations in carbon sink pathways on photosynthetic performance. Algal Research 2016;20:87–99.

Hudson GS, Evans JR, von Caemmerer S et al. Reduction of ribulose-1,5-bisphosphate carboxylase/oxygenase content by antisense RNA reduces photosynthesis in transgenic tobacco plants. Plant Physiol 1992;98:294–302.

Jiang Y-L, Wang X-P, Sun H et al. Coordinating carbon and nitrogen metabolic signaling through the cyanobacterial global repressor NdhR. Proc Natl Acad Sci U S A 2018;115:403–8.

Klähn S, Orf I, Schwarz D et al. Integrated Transcriptomic and Metabolomic Characterization of the Low-Carbon Response Using an *ndhR* Mutant of *Synechocystis* sp. PCC 6803. Plant Physiol 2015;169:1540–56.

Kopf M, Klähn S, Scholz I et al. Comparative analysis of the primary transcriptome of *Synechocystis* sp. PCC 6803. DNA Res 2014;21:527–39.

Kraus A, Hess WR. How Small Proteins Adjust the Metabolism of Cyanobacteria Under Stress: The Role of Small Proteins in Cyanobacterial Stress Responses. Bioessays 2025;47:e202400245.

Kraus A, Spät P, Timm S et al. Protein NirP1 regulates nitrite reductase and nitrite excretion in cyanobacteria. Nat Commun 2024;15:1911.

Larkin MA, Blackshields G, Brown NP et al. Clustal W and Clustal X version 2.0. Bioinformatics 2007;23:2947–8.

Liu X-Y, Hou W-T, Wang L et al. Structures of cyanobacterial bicarbonate transporter SbtA and its complex with PII-like SbtB. Cell Discov 2021;7:63.

Long BM, Rae BD, Rolland V et al. Cyanobacterial CO_2_-concentrating mechanism components: function and prospects for plant metabolic engineering. Curr Opin Plant Biol 2016;31:1–8.

Mantovani O, Reimann V, Haffner M et al. The impact of the cyanobacterial carbon-regulator protein SbtB and of the second messengers cAMP and c-di-AMP on CO_2_-dependent gene expression. New Phytol 2022;234:1801–16.

Mitschke J, Georg J, Scholz I et al. An experimentally anchored map of transcriptional start sites in the model cyanobacterium *Synechocystis* sp. PCC6803. *Proc* Natl Acad Sci U S A 2011;108:2124–9.

Nieves-Morión M, Romero-García R, Bardi S et al. Retention of a SulP-family bicarbonate transporter in a periplasmic N_2_-fixing cyanobacterial endosymbiont of an open ocean diatom. ISME J 2025;19:wraf202.

Ogawa T. A gene homologous to the subunit-2 gene of NADH dehydrogenase is essential to inorganic carbon transport of *Synechocystis* PCC6803. Proc Natl Acad Sci U S A 1991;88:4275–9.

Omata T, Gohta S, Takahashi Y et al. Involvement of a CbbR homolog in low CO_2_-induced activation of the bicarbonate transporter operon in cyanobacteria. J Bacteriol 2001;183:1891–8.

Omata T, Price GD, Badger MR et al. Identification of an ATP-binding cassette transporter involved in bicarbonate uptake in the cyanobacterium *Synechococcus* sp. strain PCC 7942. Proc Natl Acad Sci U S A 1999;96:13571–6.

Orf I, Schwarz D, Kaplan A et al. CyAbrB2 Contributes to the Transcriptional Regulation of Low CO_2_ Acclimation in *Synechocystis* sp. PCC 6803. Plant Cell Physiol 2016;57:2232–43.

Orthwein T, Scholl J, Spät P et al. The novel PII-interactor PirC identifies phosphoglycerate mutase as key control point of carbon storage metabolism in cyanobacteria. Proc Natl Acad Sci U S A 2021;118, DOI: 10.1073/pnas.2019988118.

Price GD, Badger MR, Woodger FJ et al. Advances in understanding the cyanobacterial CO_2_-concentrating-mechanism (CCM): functional components, Ci transporters, diversity, genetic regulation and prospects for engineering into plants. J Exp Bot 2008;59:1441–61.

Price GD, Woodger FJ, Badger MR et al. Identification of a SulP-type bicarbonate transporter in marine cyanobacteria. Proc Natl Acad Sci U S A 2004;101:18228–33.

Rae BD, Förster B, Badger MR et al. The CO_2_-concentrating mechanism of *Synechococcus* WH5701 is composed of native and horizontally-acquired components. Photosynth Res 2011;109:59–72.

Raven JA, Cockell CS, De La Rocha CL. The evolution of inorganic carbon concentrating mechanisms in photosynthesis. Philos Trans R Soc Lond B Biol Sci 2008;363:2641–50.

Rippka R, Deruelles J, Waterbury JB et al. Generic assignments, strain histories and properties of pure cultures of cyanobacteria. J Gen Microbiol 1979;111:1–61.

Rourke LM, Byrt CS, Long BM et al. Functional characterisation of bicarbonate transporters from the cyanobacterial SbtA2 family and subsequent expression in tobacco. 2025:2025.11.03.686429.

Rousseaux CS, Gregg WW. Interannual Variation in Phytoplankton Primary Production at A Global Scale. Remote Sensing 2014;6:1–19.

Saschenbrecker S, Bracher A, Rao KV et al. Structure and function of RbcX, an assembly chaperone for hexadecameric Rubisco. Cell 2007;129:1189–200.

Scanlan DJ, Ostrowski M, Mazard S et al. Ecological genomics of marine picocyanobacteria. Microbiol Mol Biol Rev 2009;73:249–99.

Schuller JM, Saura P, Thiemann J et al. Redox-coupled proton pumping drives carbon concentration in the photosynthetic complex I. Nat Commun 2020;11:494.

Schwarz D, Schubert H, Georg J et al. The gene sml0013 of Synechocystis species strain PCC 6803 encodes for a novel subunit of the NAD(P)H oxidoreductase or complex I that is ubiquitously distributed among Cyanobacteria. Plant Physiol 2013;163:1191–202.

Selim KA, Haase F, Hartmann MD et al. PII-like signaling protein SbtB links cAMP sensing with cyanobacterial inorganic carbon response. Proc Natl Acad Sci U S A 2018;115:E4861–9.

Selim KA, Haffner M, Burkhardt M et al. Diurnal metabolic control in cyanobacteria requires perception of second messenger signaling molecule c-di-AMP by the carbon control protein SbtB. Sci Adv 2021;7:eabk0568.

Shibata M, Katoh H, Sonoda M et al. Genes essential to sodium-dependent bicarbonate transport in cyanobacteria: function and phylogenetic analysis. J Biol Chem 2002;277:18658–64.

Song K, Baumgartner D, Hagemann M et al. AtpΘ is an inhibitor of F0F1 ATP synthase to arrest ATP hydrolysis during low-energy conditions in cyanobacteria. Curr Biol 2022;32:136–148.e5.

Spät P, Krauspe V, Hess WR et al. Deep Proteogenomics of a Photosynthetic Cyanobacterium. J Proteome Res 2023;22:1969–83.

Spreitzer RJ, Salvucci ME. Rubisco: structure, regulatory interactions, and possibilities for a better enzyme. Annu Rev Plant Biol 2002;53:449–75.

Steglich C, Futschik ME, Lindell D et al. The challenge of regulation in a minimal photoautotroph: non-coding RNAs in *Prochlorococcus*. PLoS Genet 2008;4:e1000173.

Storz G, Wolf YI, Ramamurthi KS. Small proteins can no longer be ignored. Annu Rev Biochem 2014;83:753–77.

Thiel T, Bramble J, Rogers S. Optimum conditions for growth of cyanobacteria on solid media. FEMS Microbiol Lett 1989;52:27–31.

Von Caemmerer S, Millgate A, Farquhar GD et al. Reduction of Ribulose-1,5-Bisphosphate Carboxylase/Oxygenase by Antisense RNA in the C4 Plant Flaveria bidentis Leads to Reduced Assimilation Rates and Increased Carbon Isotope Discrimination. Plant Physiol 1997;113:469–77.

Wang H-L, Postier BL, Burnap RL. Alterations in global patterns of gene expression in *Synechocystis* sp. PCC 6803 in response to inorganic carbon limitation and the inactivation of ndhR, a LysR family regulator. J Biol Chem 2004;279:5739–51.

Xu M, Bernát G, Singh A et al. Properties of mutants of *Synechocystis* sp. strain PCC 6803 lacking inorganic carbon sequestration systems. Plant Cell Physiol 2008;49:1672–7.

Yeates TO, Kerfeld CA, Heinhorst S et al. Protein-based organelles in bacteria: carboxysomes and related microcompartments. Nat Rev Microbiol 2008;6:681–91.

Zhang J, Gao F, Zhao J et al. NdhP is an exclusive subunit of large complex of NADPH dehydrogenase essential to stabilize the complex in *Synechocystis* sp. strain PCC 6803. J Biol Chem 2014;289:18770–81.

Zinchenko VV, Piven IV, Melnik VA et al. Vectors for the complementation analysis of cyanobacterial mutants. Russ J Genet 1999;35:228–32.

