## Supplementary Material for "The small protein SbtC is a functional component of the CO_2_ concentrating mechanism of cyanobacteria"

**Supplementary Information**

*-Supplementary Tables -*

**Supplementary Table 1: Primers used for cloning and genotyping.** Underlined sequences represent restriction endonuclease sites, grey labelling refers to terminator sequences.

| Primer name | Sequence (5′→3′) | Description |
|---|---|---|
| 1641_up_fw | CGTTGTTGAGGCTATAAGTTGAGGC | Amplification of homologous flanks for *sbtC* deletion. PCR product covers the genomic region upstream of *sbtC* |
| 1641_up_rev | agcgtagttcatatgTGCAATATTTGGTTCCTAAGCAGTATC | |
| 1641_down_fw | agcgtagttgcggccgcCGCCATAGTCCTCCGCTAGAATC | Amplification of homologous flanks for *sbtC* deletion. PCR product covers the genomic region downstream of *sbtC* |
| 1641_down_rev | TTTACTATTATTTGGCGATCGCCAGC | |
| Gm_fw_NotI | agcgtagttgcggccgcGAATTGACATAAGCCTGTTCGGTTCG | Amplification of a gentamicin resistance cassette (Gm$^R$) for the ligation with *sbtC* up- and downstream regions via NotI and NdeI. Downstream of Gm$^R$ an oop terminator sequence (highlighted in grey) was added. |
| Gm_rev_oop_NdeI | agcgtagttcatatg*aataaaaaacgcccggcggcaaccgagcgtt*CCGGGCTATGTGCAACGGGAAT | |
| 1641_seq_fw | TAGAGGTAGAATGATTCCCGCCAG | Amplification of final construct for *sbtC* deletion |
| 1641_seq_rev | ATGCAGGTTTGAATGCCTATGTCTTG | |
| Psbt_fw | GGGGCAATGCTCATAATCACC | Amplification of genomic region harboring *sbtABC* (incl. ~600 bp upstream of *sbtC* and ~550 bp downstream of *sbtB (slr1513))* |
| Psbt_rv | GGACGCCATTGTCATTACTC | |
| P1_sbtC$^{ery}$ | gggagaaagggaagaactgggc | Amplification of upstream region for *sbtC* deletion via homologous recombination in Δ5::*sbtA.* Introduction of a linker to erythromycin resistance cassette (Ery$^R$) |
| P2_sbtC$^{ery}$ | TCAGATGCGGACTCTAGAGGATCCatggtgattatgagcattgcccc | |
| P3_sbtC$^{ery}$ | ggggcaatgctcataatcaccatGGATCCTCTAGAGTCCGCATCTGA | Amplification of erythromycin resistance cassette (Ery$^R$) for the fusion to up and downstream flanks of *sbtC* |
| P4_sbtC$^{ery}$ | cccctgtaagttgatcaatgggacTCTAGAGGATCCCGCGGTACC | |
| P5_sbtC$^{ery}$ | GGTACCGCGGGATCCTCTAGAgtcccattgatcaacttacagggg | Amplification of downstream region for *sbtC* deletion via homologous recombination in Δ5::*sbtA.* Introduction of a linker to erythromycin resistance cassette (Ery$^R$) |
| P6_sbtC$^{ery}$ | gtcgtcaaacctaagatttcgtgcg | |
| PpetE-XhoI_fw | ACTCGAGGAAGGGATAGCAAGC | Re-amplification of the P*petE*::*sbtC*::3xFLAG construct obtained by gene synthesis to add restriction endonuclease sites XhoI and HindIII for the introduction into plasmid pVZ321 |
| Toop-HindIII_rev | CAAGCTTAATAAAAAACGCCCGGC | |
| pr_sbtA_fw | CAATCCCCATAAATTTCAACCAAGGAGAC | Amplification of DNA template for the generation of single stranded RNA probes against *sbtA* |

| | | |
|---|---|---|
| pr_sbtA_rev | TAATACGACTCACTATAGGGAGAGCCAAGGGCGGCAATAACCATC | |
| pr_1641_fw | CACCATAGAGTGAAATCCATGAACAAG | Amplification of DNA template for the generation of single stranded RNA probes against *sbtC* |
| pr_1641_rev | TAATACGACTCACTATAGGGAGAGCCAAGCGTGGGAAGGTTC | |

**Supplementary Table 2: *Synechocystis* mutant strains generated and used in this studyand their respective antibiotic resistances.** Gm – gentamicin, Hyg – hygromycin, Cm – chloramphenicol, Km – kanamycin, Sp – spectinomycin, Ery – erythromycin.

| **Strain** | **Selection markers** | **Strain description** | **Source** |
|---|---|---|---|
| WT | - | *Synechocystis* sp. PCC 6803 substrain Kazusa (Wildtype) | Norio Murata, Kazusa institute, Japan |
| Δ*sbtC* | Gm | *Synechocystis* strain with a deletion of the *sbtC* gene, achieved by replacement with gentamicin resistance cassette | This study |
| Δ5 | Cm, Hyg, Gm, Km, Sp | *Synechocystis* strain in which all Ci uptake systems are deleted, Genotype: *sbtA*::Cm, *cmpA*::Hyg, *bicA*::Gm, *ndhD4*::Km, *ndhD3*::Sp | (Xu *et al.* 2008) |
| Δ5::*sbtA* | Hyg, Gm, Km, Sp | Δ5 strain in which *sbtA* has been restored, Genotype: *cmpA*::Hyg, *bicA*::Gm, *ndhD4*::Km, *ndhD3*::Sp | This study |
| Δ5::*sbtA* Δ*sbtB* | Hyg, Gm, Km, Sp, Ery | Δ5::*sbtA* in which *sbtB* was separately deleted, Genotype: *cmpA*::Hyg, *bicA*::Gm, *ndhD4*::Km, *ndhD3*::Sp, *sbtB*::Ery | This study |
| Δ5::*sbtA* Δ*sbtC* | Hyg, Gm, Km, Sp, Ery | Δ5::*sbtA* in which *sbtC* was separately knocked out, Genotype: *cmpA*::Hyg, *bicA*::Gm, *ndhD4*::Km, *ndhD3*::Sp, *sbtC*::Ery | This study |
| WT+pVZ321 | Km, Cm | WT strain harboring an empty pVZ321 plasmid | This study |
| WT+ pVZ321-P*petE*::*sbtC*::3xFLAG | Km, Cm | WT strain harboring a pVZ321 plasmid with introduced construct P*petE*::*sbtC*::3xFLAG | This study |
| Δ*sbtC*+pVZ321 | Km, Cm, Gm | Δ*sbtC* strain harboring an empty pVZ321 plasmid | This study |
| Δ*sbtC*+pVZ321-P*petE*::*sbtC*::3xFLAG | Km, Cm, Gm | Δ*sbtC* strain harboring a pVZ321 plasmid with introduced construct P*petE*::*sbtC*::3xFLAG | This study |

*-Supplementary Figures -*

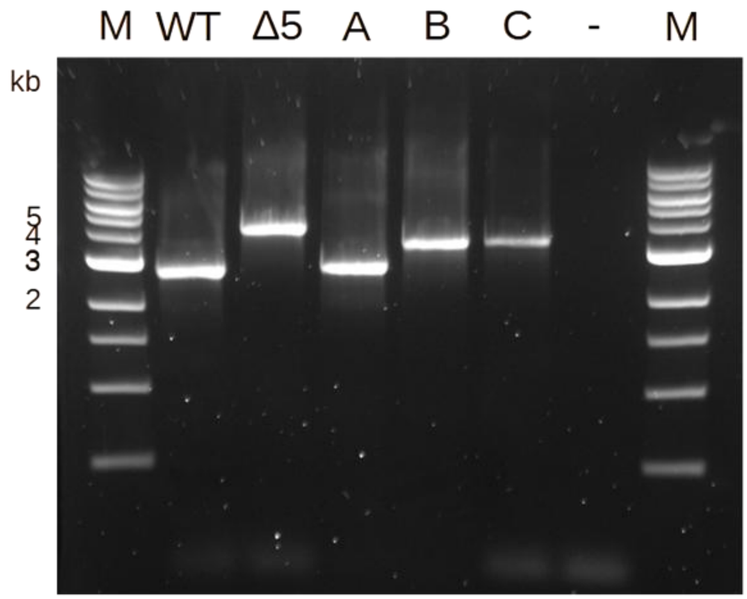


**Supplementary Figure S1: Genotyping the *sbtA* region.** M - 1kb ladder (NEB), WT - wild type, Δ5, A - Δ5::*sbtA,* B - Δ5::*sbtA*Δ*sbtB*$^{ery}$, C - Δ5::*sbtA*Δ*sbtC*$^{ery}$, – negative control. Primers used ($P_{fw}$ and $P_{rv}$) are listed in Supplementary Table 1.

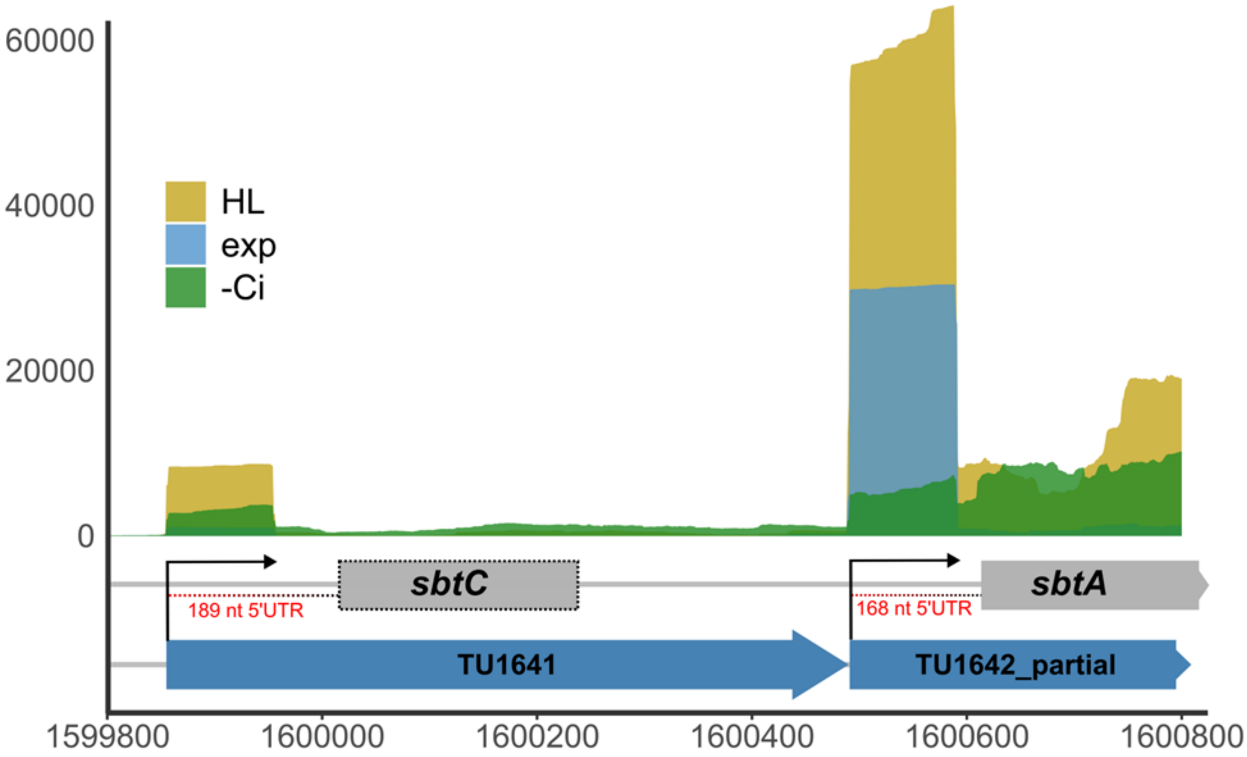


**Supplementary Figure S2: Localization of *sbtC* upstream of the *sbtA* gene and differential RNA-seq coverage for both genes under three different conditions (data replotted from** (Kopf *et al.* 2014)**.** These conditions were: –Ci, cells were grown at ambient $CO_2$ in standard BG11 medium, washed three times with $C_i$-free BG11, and cultivation was continued for 20 h; HL, high light, 470 µmol photons $m^{-2}$ $s^{-1}$ for 30 min; exp, exponential growth phase cells. Mapped initiation sites of transcription are indicated by arrows; the respective transcriptional units (TU1641 and TU1642) and length of the respective 5′-UTRs are indicated. The positions within the chromosome are plotted on the x-axis. Values at y-axis indicate normalized sequencing read counts for the primary 5′ ends x50. The positions match to the *Synechocystis* chromosome sequence in Genbank accession NC_000911. This figure extends **Fig. 1** and **Fig. 2**.

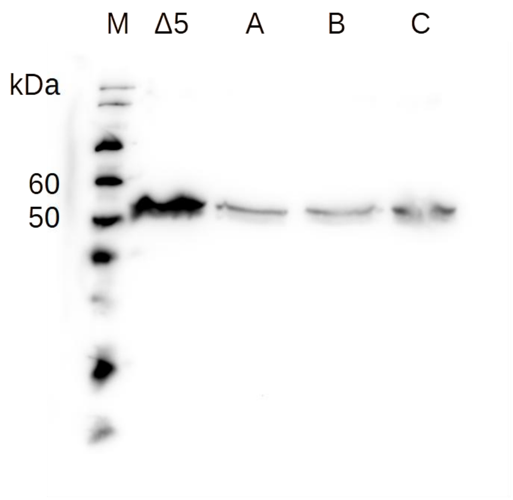


**Supplementary Figure S3: RbcL immunodetection in protein extracts with equal amounts of 1.9 µg Chl a from the strains used in the experiments shown in Fig. 4.** M – Marker, WesternFroxx (NeoFroxx) protein ladder, ∆5, A - Δ5::*sbtA,* B - Δ5::*sbtA*Δ*sbtB*ery, C - Δ5::*sbtA*Δ*sbtC*ery. RbcL/Chl a ratio is highly increased in ∆5 and seems to be slightly increased in Δ5::*sbtA*Δ*sbtC*ery.

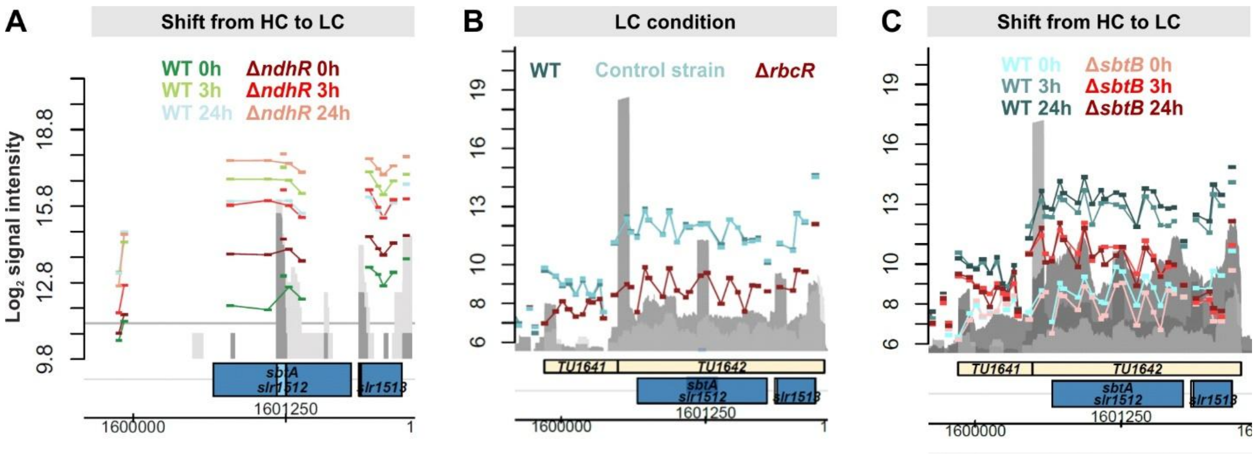


**Supplementary Figure S4: Expression of *sbtC* and *sbtAB* in various *Synechocystis* mutants affected in Ci-dependent signal cascades.** The data were extracted from previously performed microarrays and particularly show the genomic region encoding SbtA, B and C. **A:** LC-induced expression in WT and Δ*ndhR* (data from (Klähn *et al.* 2015)). Deletion of *ndhR* results in the partial de-repression of *sbtAB* pointing at its role as repressor. In contrast, *sbtC* (=TU1641, covered by the probes upstream of *sbtAB*) is not affected under HC. However, the final LC expression level of *sbtAB* is still higher, which points at an additional positive regulator under LC. **B:** Transcript abundance in a partial knockout strain for the essential transcriptional regulator RbcR under consecutive LC conditions (data extracted from (Bolay *et al.* 2022)). Apparently, full expression of both *sbtAB* and *sbtC* requires sufficient amounts of RbcR. **C:** LC-induced expression in Δ*sbtB* (data from (Mantovani *et al.* 2022)). Since, the LC-level of both *sbtA* and *sbtC* is also lower in cells lacking SbtB, this PII-like protein could be somehow involved in the positive regulation of these genes as well.

*-Supplementary Sequences-*

**Genetic construct for $Cu^{2+}$-inducible expression of *sbtC* generated by gene synthesis**

NNN – PpetE
NNN – ORF
NNN – 3xFLAG
NNN – UTRs
NNN – Toop
TAG – change of AT to TA and C to G to destroy putative NdhR binding motif
**A** – transcriptional start site (+1)
nnn –sequences containing endonuclease sites for transfer into plasmid pVZ321 (subsequently added via PCR amplification, sites included in the used primers PpetE-XhoI_fw/Toop-HindIII_rev)

>P*petE*::*sbtC*::3xFLAG::oop_XhoI/HindIII
actcgagGAAGGGATAGCAAGCTAATTTTTATGACGGCGATCGCCAAAAACAAAGAAAATTCA
GCAATTACCGTGGGTAGCAAAAAATCCCCATCTAAAGTTCAGTAAATATAGCTAGAACAACC
AAGCATTTTCGGCAAAGTACTATTCAGATAGAACGAGAAATGAGCTTGTTCTATCCGCCCGG
GGCTGAGGCTGTATAATCTACGACGGGCTGTCAAACATTGTGATACCATGGGCAGAATATA
GTTTTTTCTAAAAAATAAGTCTTATTTGTATCTATTGAATCGGGGCAATTTAAACTCAGAATA
GATTAGTTGTTCCCAGCTGAAACCATCGTGTGCTTTTTCCAGAGGCGTTTTTGGCAATTTTT
CCTCTGGTAAATTTCACCGACTTTGGGGCAATGCTCATAATCACCATAGAGTGAAATCCATG
AACAAGTTTGAATCAAGACAATCGGAACCAAAGACAAATAACAATGCTGGTTGGATAATTCA
TGTGTATGATAAAAGTCGCCGTCTTTTATTTGTTCTAGAACCTTCCCACGCTTGGCTCTTTTT
CCTGGGTTGTGGTGTCGGCTTACTACTGTCGGTGGTTTGGGTCAATGTTGCTCGCCATAGT
CCTCCGCTAGAATCCTCCCCAGTCAAGGTCTCGCCTCCCTTCCAGGTCGACGATTATAAAG
ATCATGATGGCGATTATAAAGATCATGATATTGATTATAAAGATGATGATGATAAATAGCACA
ACAATTTAAAAATCAGAAAAATTGTCCCATTGATCAACTTACAGGGGGCCATTGAGCAAAAT
CCGGGGTCACCATCTAGTCCCCAAAAAGCTGGCGATCGCCAAATAATAGTAAAACTTATCA
TTCAAATTTAAAATTACTTAGCAGATCCAGGGGGACAACTGCAAAATTGGTCGGATTTACAT
ATAGACTTTAGCTTTTCGCTCGGTTGCCGCCGGGCGTTTTTTATTaagcttg
